# Biotic interactions and climate in species distribution modelling

**DOI:** 10.1101/520320

**Authors:** Daniel P. Bebber, Sarah J. Gurr

## Abstract

Species have preferred environmental niches ^1^ and their geographical distributions respond to global climate change ^2^. Predicting range shifts under climate change has profound implications for conservation of biodiversity ^3^, provision of ecosystem services, and in the management of invasive species ^4^. Species distribution modelling (SDM) has largely focussed on climate variations, but biotic interactions, such as predation and competition, can alter potential distributions ^5,6^ and affect migration rates ^7^. However, a lack of data on biotic interactions has restricted consideration of these factors for many species ^1^. Here, we compare the power of biotic and climatic factors as predictors of global distributions of hundreds of crop pests and pathogens (CPPs), for which host preferences are known. We show that host availability is a more important predictor of endobiotic pathogen distributions (fungi, oomycetes, bacteria, viruses and nematodes) than of epibiotic pest distributions (insect herbivores). Conversely, climatic variables are better predictors of epibiotic pest distributions. These results are robust to statistical controls for varying observational capacity among countries. Our findings demonstrate that life history affects global scale species distributions and that SDM should incorporate biotic interactions as well as climate.

---

The strong influence of climate on species distributions has motivated the field of Species Distribution Modelling (SDM)^8^. In SDM, species’ climatic preferences and tolerances, derived by experiment or inferred from observed distributions, are used to project how the location of suitable habitat may change in the future ^8^. While the methods and approaches for climate-driven SDM have invoked some controversy ^9,10^, there has been little discussion of the other factors that determine where species may be found ^1,6^. Among these are various biotic interactions, such as the availability of food or interference from competitors ^1,5^, and the ability of species to reach a suitable habitat - illustrated by the rapid global redistribution of species by human activities in recent times ^11^.

The importance of biotic interactions and dispersal in determining species distributions has been distilled in the Biotic-Abiotic-Migration (BAM) framework^1,6^. Species dispersal limitation has received considerable attention, for example, certain species seem unable to keep up with the rate of climatic change ^12^. In contrast, the role of biological factors in modifying distributions is poorly understood, because the complex network of interactions that could influence species distribution is largely unknown ^1^. While examples of the roles that biotic factors can play in moderating migration rates, and thus non-equilibrium distributions, are beginning to accumulate ^7^, there remains a lack of synthesis of the general patterns and principals governing the importance of biological interactions in SDM. Soberon and Nakamura ^1^ state that “Without a much larger empirical database about such [biotic] factors, the relative roles of [abiotic] and [biotic] factors in determining distributions would be difficult to assess”.

Here, we address this question using the distributions of the pests and pathogens that attack agricultural crops. Crop pests and pathogens (CPPs) comprise a diverse group of organisms, including thousands of species and pathotypes of fungi, oomycetes, bacteria, viruses, nematodes, insects and other arthropods ^11,13^. CPPs inhabit simplified agricultural ecosystems in which intensive management practices have reduced biological diversity and physical complexity. Therefore, the network of biotic interactions in which CPPs take part is also likely to be simplified and tractable. CPPs are of great socioeconomic importance, and hence attract research interest and efforts to catalogue their biology and monitor their distributions ^11^. The distributions of many of their host plants are also known ^14^. CPPs may therefore allow the relative importance of abiotic and biotic variables in determining distributions to be quantified.

Simply finding that host availability and climatic variables are predictors of CPP presence would be uninformative because CPPs are likely to have similar climatic niches to their hosts, and therefore the two predictors will be correlated. Rather, we compared the relative predictive power of abiotic and biotic variables for CPPs that are likely to differ in the strength of their biological interaction with, and dependence upon, their hosts. Invasive endobiotic pathogens such as the viruses, bacteria, oomycetes and fungi are “buffered” to some degree from the external environment by their respective hosts. Pathogen growth rates, for example, may be better described by temperatures within host plant tissues rather than by external air temperature ^15^. Thus, their presence should be more dependent upon host distributions than are epibiotic arthropod pests, which are likely to be more directly exposed to the weather.

Generalized Additive Models (GAMs) fitted to presence/absence data explained a mean of 16.8 ± 0.3 % of the deviance using host availability, 27.8 ± 0.4 % using climatic variables, and 37.7 ± 0.5 % using both host and climate predictors. There was significant variation among CPP taxonomic groups, with biotic variables being better predictors for pathogens than for pests (Fig. 1). The difference D_diff_ in the fraction of deviance explained between climate (D_clim_) and host (D_host_) was 6.9 ± 1.5 % for pathogens, and 15.6 ± 0.9 % for pests (mixed effects model with random intercepts per taxonomic group, F_1,8_ = 17.54, p = 0.003). There was no significant influence of host number on the difference in D_diff_ between pests and pathogens (mixed effects model including log host number, F_1,1029_ = −0.49, p = 0.63).

**Fig 1.**
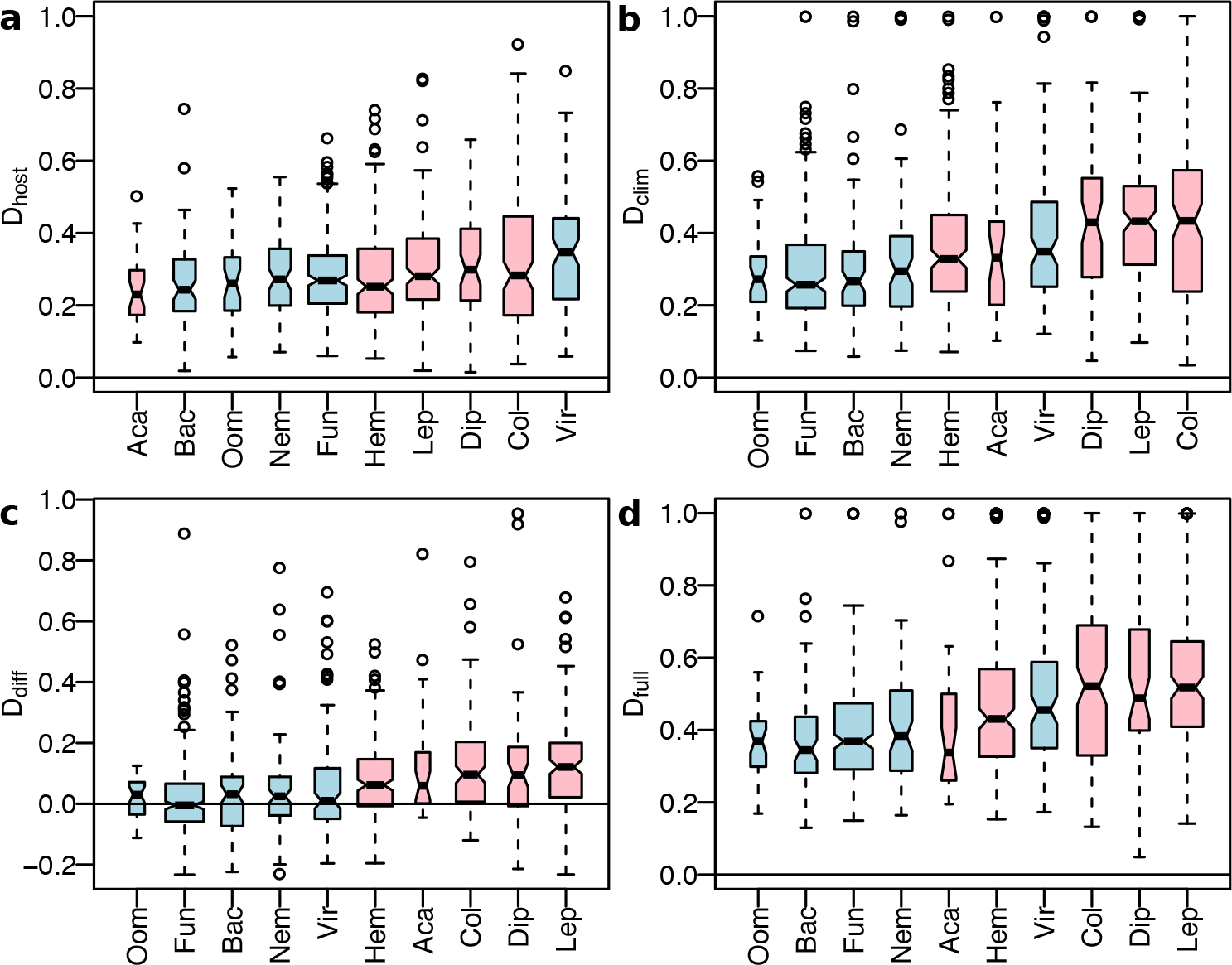
Explanatory power of GAMs. a. Deviance explained by host availability, D_host_, for each CPP category in order of mean. Pathogens are blue and pests pink. b. Deviance explained by the climate model, D_clim_. c. Difference between D_clim_ and D_host_, D_diff_,. d. Deviance explained by all predictors combined, D_full_. Where notches in the boxes do not overlap between categories, this indicates a significant difference among at the 5% level. Box widths are proportional to the square root of sample size.

D_full_ declined with the number of host plants per CPP (Fig. 2; mixed effects model with random intercepts per taxonomic group, F_1,1028_ = 34.5, p < 0.001) and was greater for pests compared with pathogens (F_1,8_ = 8.0, p = 0.022), but there was no difference between pests and pathogens in the relationship to host number (F_1,1028_ = 0.022, p = 0.88). The full model explained nearly all the deviance (D_full_ > 0.95) for 75 CPPs with restricted distributions (mean of 21.1 presences). Similar results were obtained using models for which observational bias among countries was modelled as a modification of absence data related to national scientific output ^13^ (Supplementary Fig. 1).

**Fig. 2.**
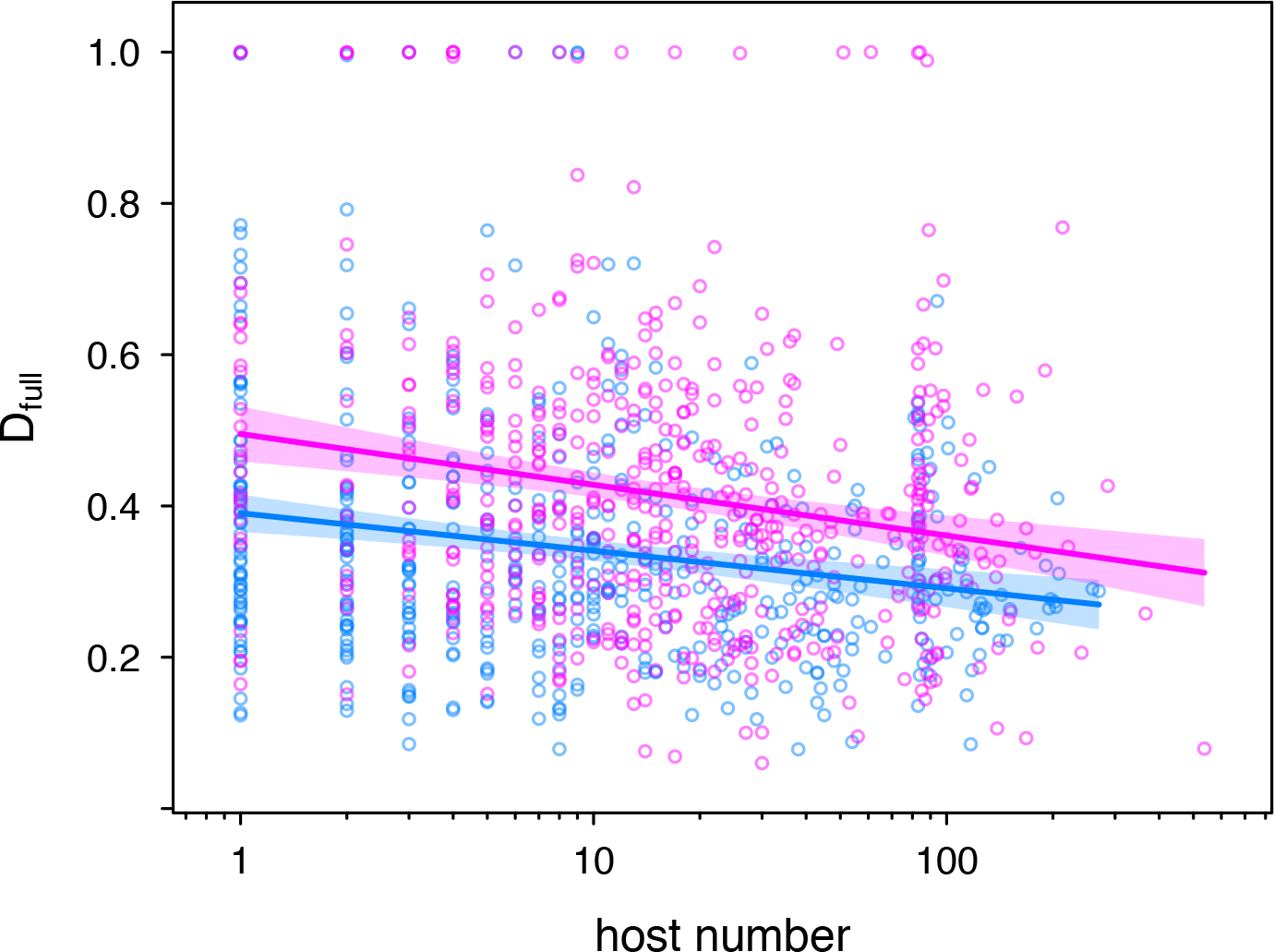
Model explanatory power declines with host range. Open circles show values for individual pests (pink) and pathogens (blue). Bold lines show linear model fits, with 95 % Confidence Limits shaded.

We found that host availability is a better predictor of species distributions for endobiotic pathogens than it is for epibiotic pests. Thus, variation in the strength of interaction between consumers and their host plants translates to biogeographic patterns at the global scale. Araújo and Luoto ^5^ first tested the hypothesis that biotic interactions can limit species distributions at macroecological scales, showing that the distribution of the clouded Apollo butterfly (*Parnassius mnemosyne*) in Europe is controlled by the distribution of host plants (*Corydalis* spp.), as well as by climate. The butterfly is threatened across much of its geographic range and has been the subject of much conservation research, hence its biotic niche has been largely described. For many other wild species such information is unavailable, and so we turned to CPPs whose biotic interactions are better described. While SDM of CPPs in relation to climate is relatively common ^16,17^, combined modelling of both pest and hosts has been rarely attempted. For example, projections of future habitat suitability for the soybean (*Glycine max*) and the bean leaf beetle (*Cerotoma trifurcata*) indicate that host distribution will limit the potential pest distribution ^18^. The soybean shows wide climatic tolerance, but the temperature range of the beetle, a generalist feeding on numerous hosts, is even wider.

Detecting any influence of host availability is perhaps surprising because we expect CPPs to be highly-adapted to their hosts and likely to evolve to match the host’s climatic niche. Therefore, the biotic and climatic predictors should be highly correlated. For example, populations of the temperate climate wheat pathogen *Zymoseptoria tritici* isolated from warmer regions show genetically-determined variation in optimal growth temperatures and temperature responses ^19^, and are therefore able to match the local climate experienced by the host. However, crops are not grown in all areas that are climatically suitable, allowing statistical models to partition the influence of these predictors. In addition, the example of the bean leaf beetle illustrates that host and pest climatic niches may not correspond exactly ^18^.

Host crops present a local environment that is substantially different from the external climate, represented by climatic variables used in SDM. Air temperature and moisture vary through a crop canopy, and leaf temperatures differ from air temperatures above the canopy depending upon insolation and leaf water status ^20^. Thus, CPPs will experience different microclimates from those suggested by meteorological data, and endobiotic pathogens, in particular, will be influenced by *in planta* conditions ^15^. SDMs utilize meteorological data and biologically relevant derivatives to estimate the climatic niche ^21,22^, but for endobiotic organisms it appears that the environment provided by the host crop is as good a predictor of presence, albeit at a global scale, as is climate. Some physiological models of pathogen risk explicitly consider the host environment. For example, a model of overwinter survival by *Phytophthora cinnamomi*, under oak bark, contains a transfer equation to calculate inner bark temperature from air temperature ^23^.

There was no significant relationship between host specificity (the number of plant genera known to be attacked by a pest or pathogen) and the relative explanatory power of biotic versus climatic predictors of species distributions. We might expect that generalist CPPs would not be limited by host availability and therefore respond more strongly to climate than specialist CPPs. However, the area of agricultural land varies widely among countries, and even a generalist able to consume any plant would therefore still be more likely found in a country with a large cropping area. Therefore, climatic predictors need not be more important for generalist CPPs. We did find that the distributions of generalist CPPs are less predictable overall, which suggests that their establishment is less hindered by the physical environment and may be determined by other factors, such as anthropogenic dispersal.

Our analysis supports the argument that biotic interactions should be included in SDM ^1,5,18^, particularly where strong dependence on other species is known or suspected. Plant viruses are obligate parasites and arguably the most strongly host-dependent of all CPPs - viral distributions appeared to be most strongly determined by host availability. However, the expected role of climate versus host availability in determining viral distributions is complicated by the fact that many plant viruses are dependent upon insect vectors, particularly aphids, for transmission ^24^. Aphids transmit nearly 300 plant viruses, and their populations respond strongly to weather ^25^. Therefore the distribution of viruses could partly reflect the climatic controls on their vectors. However, Hemiptera are the most widespread of all insect pest groups ^11^, with the cotton aphid *Aphis gossypii*, the green peach aphid *Myzus persicae*, and the silverleaf whitefly *Bemisia tabaci* most cosmopolitan. Each of these species transmits many viruses. Host crops may thus be the limiting factor in viral distributions where the vector is widespread.

We investigated only one biotic factor, host availability, which we propose is most important for CPPs. Predators of CPPs, for example, appear to be most important within the native range, their prey finding ‘enemy-free space’ upon dispersal ^27^. For wild species, many other interactions will determine the rate and pattern of range shift in response to climate change ^7^, and as empirical data accumulate, knowledge of general principles governing range shifts will improve.

The importance of weather in determining the risk of CPP outbreaks has long been recognized, resulting in numerous statistical and mechanistic models across a wide range of spatiotemporal scales ^17^. Researchers have recently begun to investigate the distributional determinants of individual CPPs and how these might develop as the climate changes. In many cases, the response of the pest is modelled, but not the host crop ^28^. This simplification could lead to serious errors in projections if, for example, the pest and host crop occupy different climatic niches ^18^. Analyses incorporating both the pest and the host are rare, and even these do not attempt to estimate the influence of dispersal limitation. We have shown that fundamental differences in life history influence the relative importance of biotic and abiotic factors in determining global distributions for a class of organisms, CPPs, for which biotic niche information is available. Accumulation of such data for wild species will help to improve SDM for projections of how the biosphere is likely to change with global warming.

## Methods

We compared the relative predictive power of abiotic (climate) and biotic (host availability) variables on current species distributions by statistically modelling the current global distributions of CPPs at national scale. We employed the approach of ‘surrogate hypotheses’ ^5^, whereby correlations in observational data are tested in lieu of controlled experiments, which are difficult to conduct for macroecological processes. We compared the goodness-of-fit of biotic (host availability) and abiotic (climate) variables to CPP presence-absence data at country level (state level for the USA, Brazil, India, China and Australia) using Generalized Additive Models (GAMs) with binomial errors ^29^. The ‘biotic model’ used the log-transformed areas of host crops for the CPP as predictors ^13^; the ‘climate model’ used the climate in areas where the host crops are grown. The ‘full model’ used both biotic and climate variables. We obtained CPP presence data, and known host plants, for each CPP from CABI with permission.

Taxonomic categories of CPPs for which fewer than twenty species were available (Psocoptera, Thysanoptera) were omitted, as were CPPs present in ten or fewer locations, leaving 1040 CPPs. We summed spatial distributions of major crops from the MIRCA2000 database ^14^ to country and state level, to match CPP distribution data. For each country/state, we calculated the total production area of each crop, and for each CPP we calculated the total production area for each host crop. Host crop areas were log-transformed for fitting. We selected four bioclimatic variables from the BIOCLIM database ^30^: mean annual temperature; standard deviation of monthly temperature; mean annual rainfall; standard deviation of monthly rainfall. We omitted the other BIOCLIM variables (annual temperature range, precipitation of driest month, etc.) to avoid multi-collinearity, as these derived variables are strongly correlated with those we selected. Area-weighted means of these climatic variables were calculated for crop-growing areas per region. GAMs were fitted using splines for each predictor variable ^31^. We analysed the difference between the fraction of deviance explained by the abiotic and biotic models to determine whether the relative importance of host availability varied between pests and pathogens.

Available global CPP distribution data are biased by the varying abilities of countries to detect, identify and report CPP presence. Richer countries at higher latitudes tend to have better plant health reporting systems, and there is a strong correlation between countries’ scientific output and reported pest incidence ^4,13^. While presences reported in the CABI databases are likely to be reliable due to stringent quality checking procedures, absences could be pseudo-absences where a CPP is present but unreported. Let *d*_*n*_ be the binary presence-absence data for a pest or pathogen in each of *n* geographical regions, and let *w*_*n*_ be the corresponding confidence weighting for that region. Then the confidence-weighted presence probabilities are

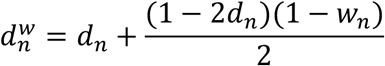

Our confidence in presences is absolute, i.e. when *d*_*n*_ = 1, *w*_*n*_ = 1

Our confidence in absences varies as a fitted quadratic function of the number of scientific publications per country between 1996 and 2012 ^13^, *s*_*n*_, such that
when *d*_*n*_ = 0, *w*_*n*_ = −0.001085 ln(*s*_*n*_ + 1) + 0.005074 ln(*s*_*n*_ + 1)^2^

Therefore, our confidence in absences from the USA is complete (the USA has the largest scientific output), *w*_*n*_ = 1, and 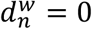 when *d*_*n*_ = 0. Our confidence in absences from the world’s least developed countries is near zero, and 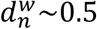. This method is not equivalent to weighting observations by confidence, nor including confidence as a predictor in the model, because the weighting only affects our treatment of absences. We fitted fourth-order polynomials of the predictor variables to 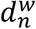, minimizing χ^2^ of the difference between observed and fitted values by optimization. Quadratic functions are often used in SDM ^29^, but quartic functions allow a greater range of response profiles to be described, without over-fitting. Presence-absence data are often fitted using Generalized Linear Models or Generalized Additive Models ^5^, but these methods require errors distributions in the exponential family for maximum likelihood estimation ^31^. The data do not follow the binomial distribution, and we are therefore unable to calculate likelihoods for fitted models, nor the various model selection metrics based upon likelihood (e.g. AIC), using GLM or GAM. Instead, we obtained χ^2^ and AUC statistics for the biotic model, the climate model, and the combined model. These values were then compared using linear models to investigate the relative explanatory power of biotic and abiotic predictors.

## Acknowledgements

Data on pest and pathogen geographical distributions and host plants were obtained with permission from CABI.

## Supplementary Information

### Supplementary Figures

**Supplementary Fig. 1.**
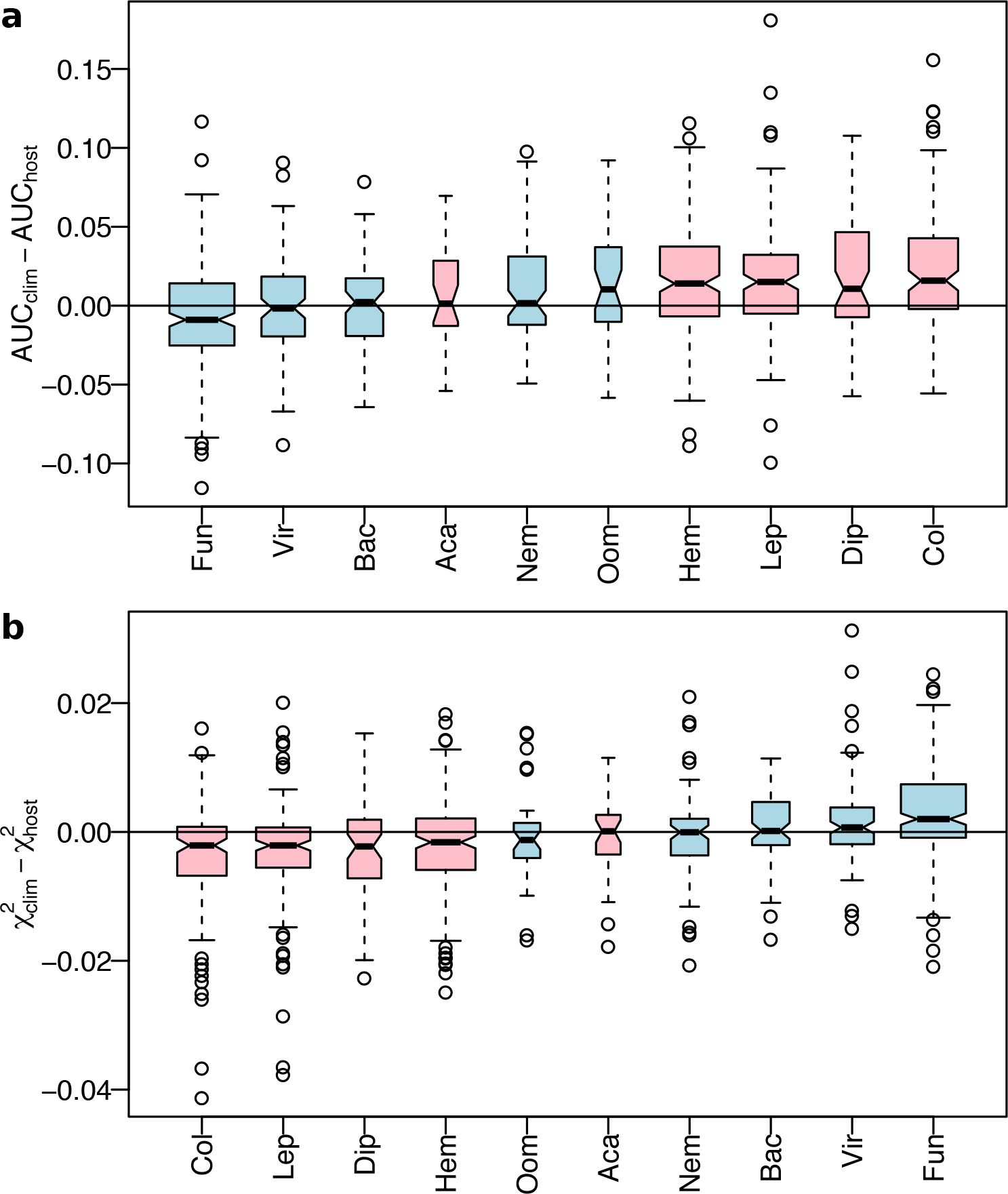
Explanatory power of climate vs. host models. a. AUC statistic for models of CPP presence-absence adjusted for observational bias, for models based on climatic variables (AUC_clim_) vs. host availability variables (AUC_host_). Pathogen groups are blue, pest groups pink. The larger the value, the better climatic variables predict CPP distribution. AUC_diff_ was greater for invertebrate pest distributions than pathogen distributions (mixed effects model with random intercepts per taxonomic group, F_1,8_ = 12.3, p = 0.008). b. Chi-squared statistic for models of CPP presence-absence adjusted for observational bias, for models based on climatic variables (χ^2^clim) vs. host availability variables (χ^2^host). χ^2^diff was lower for invertebrate pest distributions than pathogen distributions (mixed effects model with random intercepts per taxonomic group, F_1,8_ = 16.8, p = 0.004). Where notches in the boxes do not overlap between categories, this indicates a significant difference among at the 5% level. Box widths are proportional to the square root of sample size.

